# Leveraging polygenic functional enrichment to improve GWAS power

**DOI:** 10.1101/222265

**Authors:** Gleb Kichaev, Gaurav Bhatia, Po-Ru Loh, Steven Gazal, Kathryn Burch, Malika Freund, Armin Schoech, Bogdan Pasaniuc, Alkes L Price

**Author notes:** These authors jointly supervised this work (Bogdan Pasaniuc, Alkes L Price). Correspondence should be addressed to G.K., B.P. or A.L.P.

## Abstract

Functional genomics data has the potential to increase GWAS power by identifying SNPs that have a higher prior probability of association. Here, we introduce a method that leverages polygenic functional enrichment to incorporate coding, conserved, regulatory and LD-related genomic annotations into association analyses. We show via simulations with real genotypes that the method, Functionally Informed Novel Discovery Of Risk loci (FINDOR), correctly controls the false-positive rate at null loci and attains a 9–38% increase in the number of independent associations detected at causal loci, depending on trait polygenicity and sample size. We applied FINDOR to 27 independent complex traits and diseases from the interim UK Biobank release (average *N*=130K). Averaged across traits, we attained a 13% increase in genome-wide significant loci detected (including a 20% increase for disease traits) compared to un-weighted raw p-values that do not use functional data. We replicated the novel loci in independent UK Biobank and non-UK Biobank data, yielding a highly statistically significant replication slope (0.66–0.69) in each case. Finally, we applied FINDOR to the full UK Biobank release (average *N*=416K), attaining smaller relative improvements (consistent with simulations) but larger absolute improvements, detecting an additional 583 GWAS loci. In conclusion, leveraging functional enrichment using our method robustly increases GWAS power.

## Introduction

Genome-wide Association Studies (GWAS) are the prevailing approach for identifying disease risk loci^1,2^, but the large number of statistical tests performed necessitates stringent p-value thresholds that can limit power. Emerging functional genomics data has revealed that certain categories of variants are enriched for disease heritability^3–12^. Thus, incorporating functional information into association analyses has the potential to increase GWAS power^13–21^. However, previous integrative methods for GWAS hypothesis testing either assume sparse genetic architectures when estimating functional enrichment^17,20^, require knowledge or approximation of the true effect size distribution^13–15^, or do not produce p-values for each SNP as output^17–19,21^. In addition, general-purpose methodologies for association testing that can integrate prior information^22–24^ have not been thoroughly evaluated in the context of GWAS leveraging functional genomics data.

In this work, we propose an approach that uses polygenic modeling to weight SNPs according to how well they tag functional categories that are enriched for heritability. Our procedure takes as input summary association statistics along with pre-specified functional annotations (which can be overlapping and/or continuous-valued), and outputs well-calibrated p-values. We utilize a broad set of 75 coding, conserved, regulatory and LD-related annotations that have previously been shown to be enriched for disease heritability^8,12^. We incorporate the weights computed by our method using the weighted-Bonferroni procedure described by ref. ^13^, a theoretically sound approach that ensures proper null calibration and can improve power when employed with informative weights. Through extensive simulations and analysis of UK Biobank phenotypes^25–27^, we demonstrate that our approach reproducibly identifies novel GWAS loci while controlling false positives.

## Results

### Overview of Methods

We propose an integrative GWAS framework for Functionally-Informed Novel Discovery of Risk loci (FINDOR). Our approach involves two steps. First, we use stratified LD score regression^8^ to compute the expected χ^2^ statistic of each SNP based on the functional annotations that it tags; we make use of a broad set of coding, conserved and regulatory annotations^8^ as well LD-dependent annotations^12^ (conditional on MAF, variants with lower LD have larger causal effect sizes). Second, we stratify SNPs into bins of expected χ^2^ and estimate the proportion of null 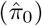 and alternative 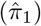 SNPs within each bin using the Storey π_0_ estimator^28^ to obtain bin-specific weights. We limit the number of bins to 100 and normalize the weights to have mean 1, ensuring proper null calibration^13^. We then divide the observed p-values within each bin by these weights to produce re-weighted p-values for each SNP. Bins with larger values of 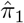 will have larger weights, leading to more significant p-values. Details of the method are described in the Online Methods section; we have released open-source software implementing the method (see URLs).

### Simulations assessing calibration and power

We assessed calibration and power via simulations using real genotypes from the UK Biobank interim release ^25^ (*N* = 100*K* subsampled British-ancestry samples, *M* = 9.6*M* well-imputed SNPs; see Online Methods). We simulated polygenic traits with 10,000 or 20,000 causal variants and SNP-heritability 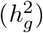 equal to 0.1 or 0.2. All causal variants were placed on odd chromosomes, with functional enrichment based on a meta-analysis of 31 traits using the baselineLD model described in ref.^12^ (Supplementary Table 1; see URLs), and even chromosomes served as null data. Weights were computed by running stratified LD score regression^8^ on association statistics computed from simulated phenotypes, without knowledge of the true functional enrichment parameters used to generate the phenotypes. We compared FINDOR to three other methods that can incorporate auxiliary information for each SNP: Stratified False Discovery Rate (S-FDR)^22^, Grouped Benjamini Hochberg (GBH)^23^, and Independent Hypothesis Weighting (IHW)^24^. For each of the four methods, we considered four different criteria for stratifying SNPs into bins: predicted χ^2^ statistics under the baselineLD model (baseLD); predicted χ^2^ statistic under the baselineLD model trained using off-chromosome data via a Leave-One-Chromosome-Out approach (baseLD-LOCO); total LD score of a SNP (LDscore), motivated by a previous study reporting that simple LD information can be used to improve GWAS power^14^; and randomly chosen bins (Random). We also considered unweighted raw p-values (Unweighted), a natural benchmark. For both null (even) and causal (odd) chromosomes, the primary metric was the number of independent genome-wide significant associations identified. Throughout this work, we define an independent association as a SNP that exceeds a significance threshold (e.g., 5 × 10^−8^), together with all linked SNPs that have an *r*^2^ > 0.01 within 5Mb. We performed 1,000 simulations and averaged results across simulations. Further details of the simulation framework are provided in the Online Methods section.

We first assessed calibration on null chromosomes. We determined that FINDOR was well-calibrated, producing a similar number of false-positive (independent, genome-wide significant) associations at null loci as the Unweighted approach (see Figure 1 and Supplementary Table 2). This remains true whether we infer functional enrichment and compute expected χ^2^ statistics using all GWAS data (baseLD) or using off-chromosome data (baseLD-LOCO), motivating the use of the baseLD stratification criteria in the remainder of this work. Similarly, FINDOR was well-calibrated at less stringent significance thresholds (see Supplementary Table 3). Although FINDOR makes multiple passes over the data, which in principle could overfit the data and produce false positives, this does not occur in practice, likely due to the small number of global parameters estimated (hundred) relative to the large number of hypothesis tests performed (millions).

**Figure 1:**
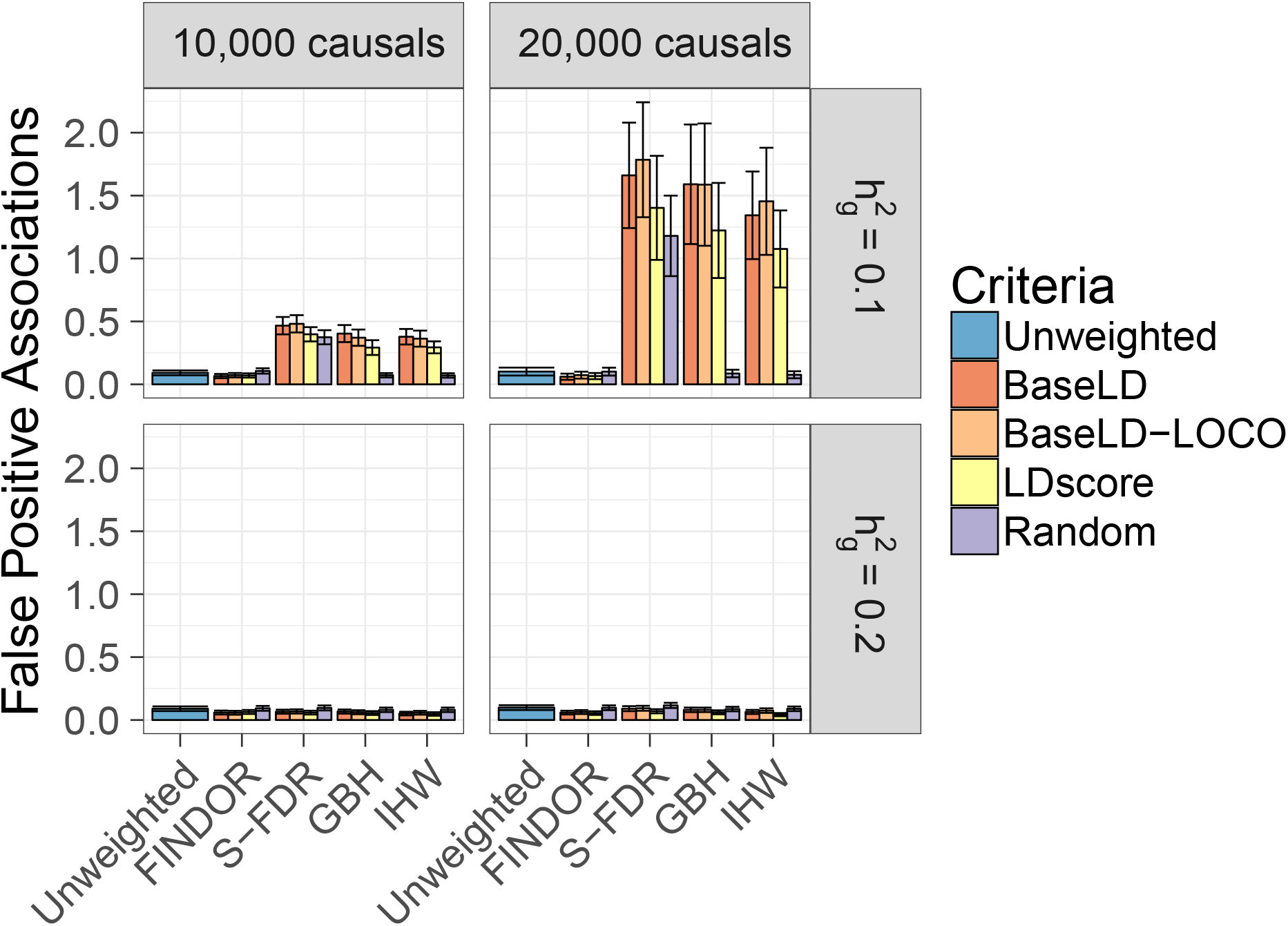
FINDOR is well-calibrated in simulations of null loci. We report the average number of independent, genome-wide significant (p < 5 × 10^−8^) associations on null chromosomes. Results are averaged across 1000 simulations. Error bars represent 95% confidence intervals. Numerical results are reported in Supplementary Table 2.

On the other hand, S-FDR, GBH and IHW each exhibited moderate to severe increases in false-positive associations, particularly at higher polygenicity and lower SNP-heritability. For example, at a polygenicity of 20,000 causal variants and 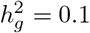, we observe an average (SE) of 0.10 (0.02) false positives per simulated GWAS using raw unweighted p-values and 0.06 (0.01) using FINDOR with baseLD criteria, while S-FDR, GBH, and IHW with baseLD yield 1.6 (0.2), 1.6 (0.2), and 1.3 (0.2) false positives, respectively (see Figure 1 and Supplementary Table 2). This inflation is exacerbated at smaller sample sizes (see Supplementary Figure 1). We hypothesize that this may be due to the fact that the theoretical guarantees provided by these procedures are unlikely to be valid when the auxiliary information incorporates the dependence structure between hypothesis tests; this limitation was previously noted by Ignatiadis et al.^24^ and clearly affects both baseLD and LDscore stratifying criteria. Furthermore, while GBH and IHW were consistently well-calibrated under random stratification (see Figure 1, purple bars), S-FDR was not, perhaps because S-FDR requires additional adjustments for the number of strata used^29^.

We next evaluated power to detect true associations on causal chromosomes. We restricted our assessment of power to Unweighted and FINDOR, as they were the only methods that were well-calibrated under the null for all stratification criteria. FINDOR attained an 8.6–38% increase in the number of true (independent, genome-wide significant) associations, depending on polygenicity (10,000 or 20,000 causal variants) and SNP-heritability (0.1 or 0.2) (see Figure 2 and Supplementary Table 4). The relative improvement was smaller at lower polygenicity and larger SNP-heritability, each of which correspond to higher absolute power. Our method has a fixed budget of weights that it can allocate, and we hypothesize that when absolute power is high it is more likely to allocate weights to SNPs that are already genome-wide significant, explaining the smaller relative improvement. In addition, the enrichment estimates provided by stratified LD score regression are expected to be less precise at lower polygenicity. However, the smaller relative improvement still translated into a larger absolute improvement in settings with higher absolute power.

**Figure 2:**
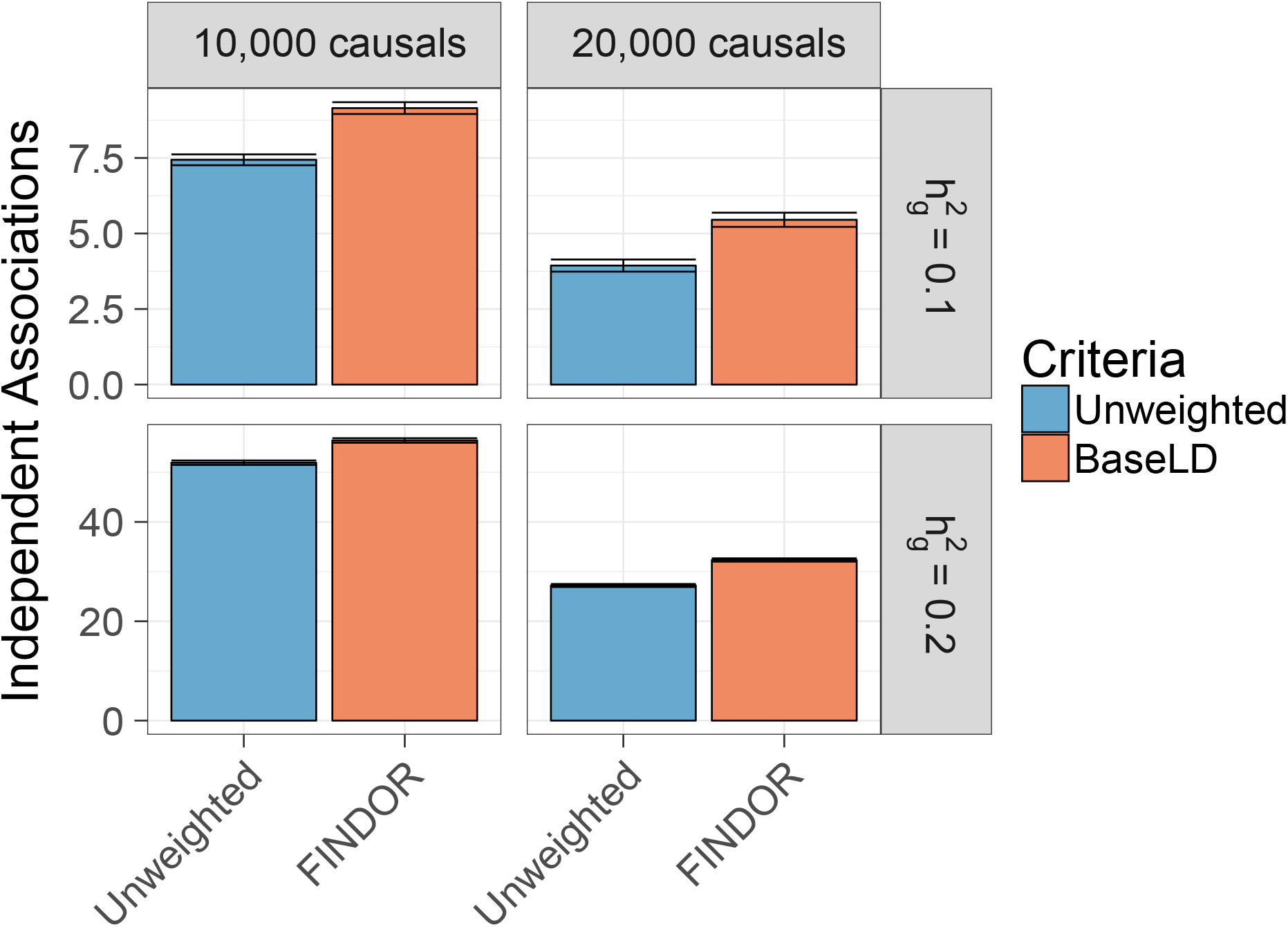
FINDOR increases power in simulations of causal loci. We report the average number of independent, genome-wide significant (*p* < 5 × 10^−8^) associations on causal chromosomes. Results are averaged across 1000 simulations. Error bars represent 95% confidence intervals. Numerical results are reported in Supplementary Table 4.

### Application to 27 UK Biobank traits

We applied FINDOR to the interim UKBiobank release^25^, which includes *N* =145*K* European-ancestry samples and *M* = 9.6*M* well-imputed SNPs. We analyzed 27 independent, highly heritable traits (average *N* =130K; see Table 1 and Online Methods). We computed summary association statistics using BOLT-LMM v2.130 (Unweighted approach). We applied FINDOR to these summary statistics and compared the number of independent, genome-wide significant associations identified by FINDOR vs. the Unweighted approach. In total, FINDOR identified 207 more associations (see Table 1 and Supplementary Tables 5 and 6), a statistically significant improvement (block-jacknife SE = 20.4, *p* < 1 × 10^−20^). This corresponds to an average per-trait improvement of 13% (SE=2.5%) and an aggregate improvement of 6.8%; FINDOR identified more associations than the Unweighted approach for 24 out of 27 traits, and the same number of associations for the remaining three traits. The aggregate improvement was lower than the average per-trait improvement because the relative improvement was smaller for traits with higher power (i.e. more associations) (see Figure 3), consistent with simulations. In particular, disease traits exhibited a larger improvement (20% average per-trait, 22% aggregate, see Supplementary Table 7), consistent with smaller effective sample size (i.e. smaller value of sample size * observed-scale SNP-heritability) due to the relatively small number of disease cases. Qualitatively similar results were obtained at a more stringent p-value threshold of 5 × 10^−9^ (see Supplementary Table 8). We note that, compared to the 13% average per-trait improvement of FINDOR with the baselineLD model^12^, FINDOR with the baseline model^8^ (which excludes LD-related annotations) attained only a 7.1% average per-trait improvement and 4.3% aggregate improvement (72 fewer GWAS hits; jackknife SE on difference = 13.3, *p* = 6.3 × 10^−8^, see Supplementary Table 5). This indicates that the LD-related annotations of the baselineLD model contain valuable information for increasing association power; in particular, these annotations avoid the phenomenon of strong LD between in-annotation and out-annotation SNPs that may limit the potential of coding, conserved and regulatory annotations to increase association power despite their strong enrichments for trait heritability.

**Figure 3:**
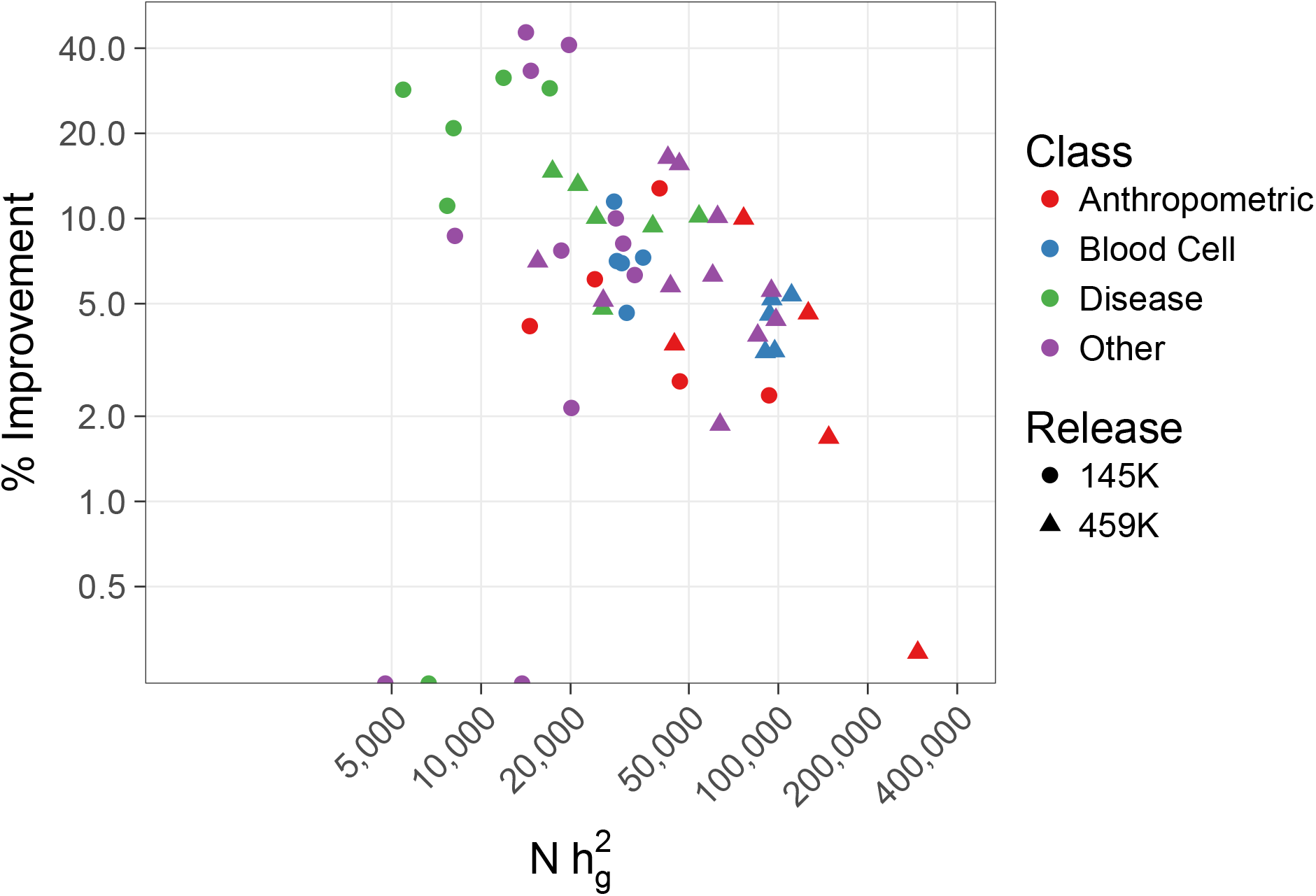
Relative improvement of FINDOR in real UK Biobank phenotypes decreases as a function of absolute power. We plot the relative improvement in the number of independent GWAS loci identified by FINDOR compared to Unweighted p-values vs. sample size times observed-scale SNP-heritability, using log scales. The three circles at the bottom of plot correspond to traits where the number of loci was identical for FINDOR compared to Unweighted p-values (0% improvement). Numerical results are reported in Table 1 and Supplementary Tables 5,6, and 12.

**Table 1:** FINDOR increases power across 27 UK Biobank traits. For each trait, we report the number of independent, genome-wide significant loci identified by the Unweighted approach and by FINDOR in the 145K and 459K UK Biobank releases. Complete results are reported in Supplementary Table 6.

| Class | Trait | N | $h_g^2$ | 145K | | | 459K | | |
| --- | --- | --- | --- | --- | --- | --- | --- | --- | --- |
|  |  |  |  | Unweighted | FINDOR | %Improve | Unweighted | FINDOR | %Improve |
| Anthropometric | Balding Type I | 68K/208K | 0.21 | 96 | 100 | 4.2% | 334 | 346 | 3.6% |
|  | Body Mass Index | 145K/458K | 0.28 | 117 | 132 | 12.8% | 908 | 950 | 4.6% |
|  | Heel T Score | 141K/446K | 0.33 | 300 | 308 | 2.7% | 1130 | 1149 | 1.7% |
|  | Height | 145K/458K | 0.64 | 674 | 690 | 2.4% | 2395 | 2402 | 0.3% |
|  | Waist-hip Ratio | 145K/458K | 0.17 | 98 | 104 | 6.1% | 460 | 506 | 10.0% |
| Blood Cell | Eosinophil Count | 140K/440K | 0.21 | 187 | 200 | 7.0% | 699 | 731 | 4.6% |
|  | Mean Corpular Hemoglobin | 141K/443K | 0.22 | 237 | 248 | 4.6% | 765 | 791 | 3.4% |
|  | Red Blood Cell (RBC) Count | 141K/445K | 0.25 | 192 | 206 | 7.3% | 840 | 885 | 5.4% |
|  | RBC Distribution Width | 141K/445K | 0.20 | 198 | 212 | 7.1% | 652 | 674 | 3.4% |
|  | White Blood Cell Count | 131K/444K | 0.21 | 148 | 165 | 11.5% | 713 | 750 | 5.2% |
| Disease | Auto Immune Traits | 145K/459K | 0.04 | 14 | 18 | 28.6% | 75 | 86 | 14.7% |
|  | Cardiovascular Diseases | 145K/459K | 0.12 | 38 | 49 | 28.9% | 285 | 314 | 10.2% |
|  | Eczema | 145K/459K | 0.08 | 35 | 46 | 31.4% | 181 | 198 | 9.4% |
|  | Hypothyroidism | 145K/459K | 0.05 | 27 | 30 | 11.1% | 139 | 153 | 10.1% |
|  | Respiratory Diseases | 145K/459K | 0.06 | 24 | 29 | 20.8% | 104 | 109 | 4.8% |
|  | Type 2 Diabetes | 145K/459K | 0.05 | 14 | 14 | 0.0% | 76 | 86 | 13.2% |
| Other | Age at Menarche | 75K/242K | 0.25 | 52 | 56 | 7.7% | 318 | 338 | 6.3% |
|  | Age at Menopause | 44K/143K | 0.11 | 18 | 18 | 0.0% | 85 | 91 | 7.1% |
|  | FEV1-FVC Ratio | 124K/370K | 0.27 | 174 | 185 | 6.3% | 684 | 714 | 4.4% |
|  | Forced Vital Capacity (FVC) | 124K/372K | 0.23 | 90 | 99 | 10.0% | 544 | 565 | 3.9% |
|  | Hair Color | 143K/452K | 0.14 | 140 | 143 | 2.1% | 428 | 436 | 1.9% |
|  | Morning Person | 130K/410K | 0.11 | 14 | 14 | 0.0% | 156 | 165 | 5.8% |
|  | Neuroticism | 124K/372K | 0.11 | 11 | 16 | 45.5% | 128 | 149 | 16.4% |
|  | Smoking Status | 145K/458K | 0.10 | 18 | 24 | 33.3% | 154 | 178 | 15.6% |
|  | Sunburn Occasion | 109K/344K | 0.07 | 23 | 25 | 8.7% | 78 | 82 | 5.1% |
|  | Systolic Blood Pressure | 134K/422K | 0.22 | 98 | 106 | 8.2% | 666 | 703 | 5.6% |
|  | Years of Education | 144K/455K | 0.14 | 17 | 24 | 41.2% | 286 | 315 | 10.1% |
|  | Overall | 145K/459K | NA | 3054 | 3261 | 6.8% | 13283 | 13866 | 4.4% |
|  | Average Per-Trait | 130K/409K | 0.18 | 113 | 120 | 13% | 491 | 513 | 6.9% |

Next, we carried out a UK Biobank-based replication analysis for the 27 traits using non-overlapping samples in the full UK Biobank release. Starting with the 459K European-ancestry samples, we excluded the 145K samples that were present in the interim release and computed summary statistics using BOLT-LMM v2.3, a highly computationally efficient implementation for very large data sets^27^. This produced a well-powered replication data set (average *N*=283K). We evaluated strength of replication by computing the replication slope, defined as the slope of a regression of estimated standardized effect sizes in replication data vs. discovery data, restricting to lead SNPs at genome-wide significant loci from the discovery data (we excluded lead SNPs that were not present in the replication data). We computed replication slopes for three classes of loci: (1) those that were genome-wide significant only using the Unweighted approach, (2) only using FINDOR p-values, or (3) using both methods. The 49 loci that were significant only using the Unweighted approach produced a replication slope of 0.57 (SE=0.043). The 230 loci that were significant only using FINDOR (i.e. novel discoveries) produced a slightly stronger replication slope of 0.66 (SE=0.018); the difference was not statistically significant based on the small number of data points, particularly for Unweighted only. As expected, the 2766 loci that were significant using both methods produced the strongest replication slope of 0.91 (SE= 0.003), as this class of loci included the most significant associations (see Figure 4 and Supplementary Table 9). We also performed a separate replication analysis for nine traits for which summary statistics from independent, non-UK Biobank GWAS were available (see Online Methods, Supplementary Table 10). In this analysis, the 36 loci that were significant only using FINDOR (i.e. novel discoveries) produced a replication slope of 0.69 (SE=0.11) in non-UK Biobank data, which did not differ significantly from the replication slope for the 410 loci that were significant using both methods (0.66, SE=0.012, see Figure 4 and Supplementary Table 11). Only a single locus was significant only using Unweighted p-values in this analysis, therefore we do not report a replication slope for this class. Overall, these results confirm that the novel loci identified by FINDOR robustly replicate in independent samples.

**Figure 4:**
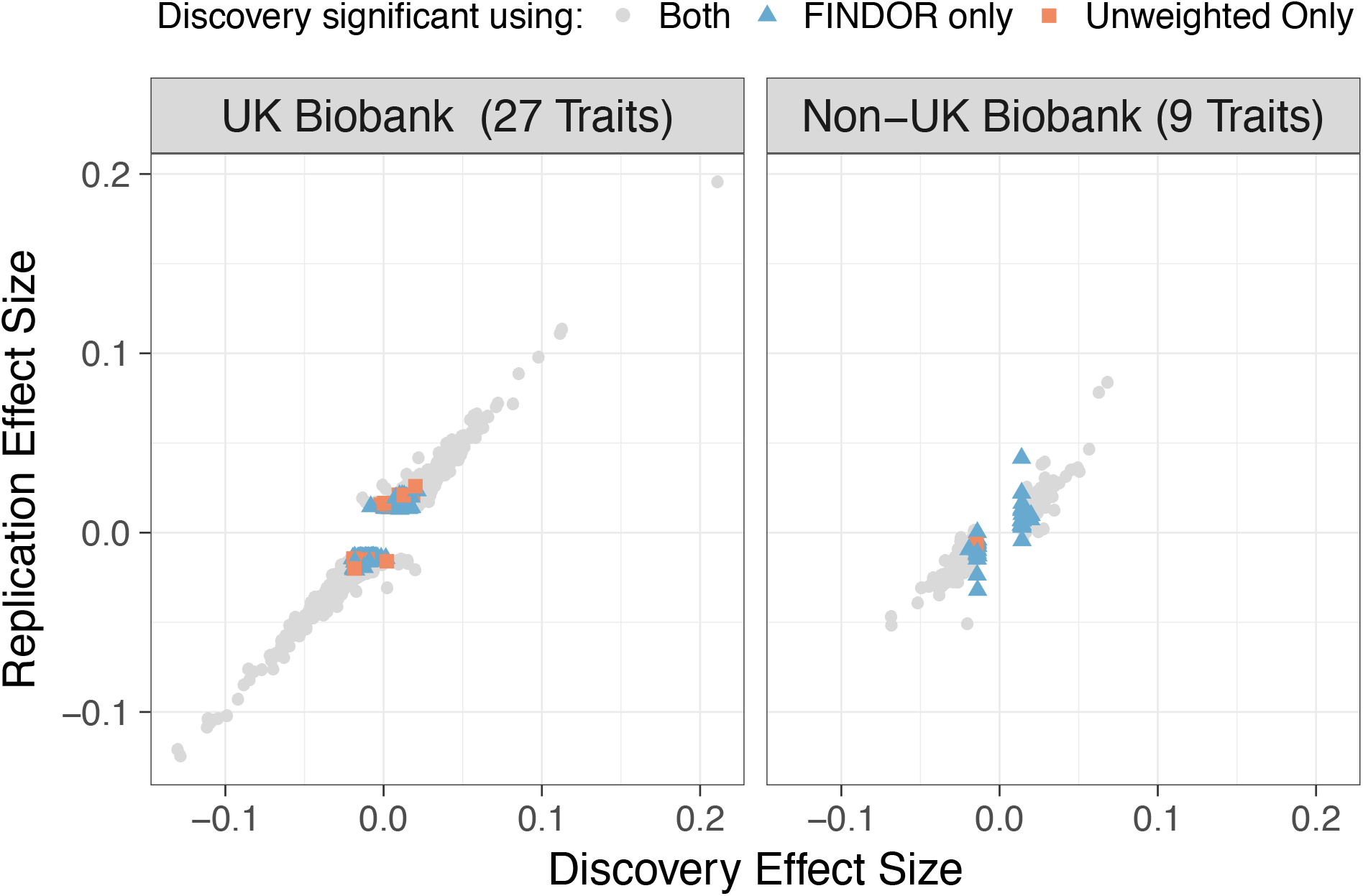
Novel loci identified by FINDOR replicate in independent samples. We plot the standardized effect sizes 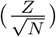 in the UK Biobank replication sample (average *N* = 283K, left panel) and non-UK Biobank replications sample (average *N* = 158K, right panel) vs. the UK Biobank discovery sample (average *N* = 132K). For novel loci identified by FINDOR (blue triangles), the replication slope was positive and highly significant in both cases (UK Biobank = 0.66, Non-UK Biobank = 0.69). Numerical results are reported in Supplementary Tables 9 and 11

Finally, we applied FINDOR to the 27 traits using the full set of 459K European-ancestry samples (average *N*=416K), analyzing summary statistics computed using BOLT-LMM v2.327. The Unweighted approach identified 13,283 independent genome-wide significant associations in this data. FINDOR identified 583 more associations (see Supplementary Table 12, Jackknife SE = 40.6, *p* < 1 × 10^−20^), corresponding to an average per-trait improvement of 6.9% (SE = 0.66%) and an aggregate improvement of 4.1% (see Table 1); FINDOR identified more associations than the Unweighted approach for all 27 traits. Once again, the relative improvements decreased as a function of sample size times observed-scale SNP-heritability (see Figure 3, Table 1), with larger relative improvements for disease traits (10% average per-trait, 10% aggregate) and smaller relative improvements in the 459K release vs. the 145K release, consistent with simulations. We further characterized Unweighted-only and FINDOR-only loci by contrasting their overlap with molecular QTL 95% causal sets^31^ (which are weakly correlated with the baselineLD model annotations used by FINDOR: |*r*| ≈ 0.05 for most annotations, see ref.^31^). The lead SNPs at FINDOR-only loci had substantial overlap with molecular QTL 95% causal sets (and substantially larger molecular QTL causal posterior probabilities on average), compared to Unweighted-only loci (see Supplementary Table 13); this implies that loci identified by FINDOR are not only more numerous, but also more amenable to biological interpretation and mechanistic insights. Overall, these results indicate that FINDOR can provide a substantial increase in power – particularly for studies with smaller effective sample sizes, such as studies of disease traits.

## Discussion

We have introduced a p-value weighting approach that leverages polygenic functional enrichment to improve association power. We demonstrated in simulations that our FINDOR framework is properly calibrated under the null and improves power to detect causal loci. We reproducibly identified hundreds of new loci across a broad set of UK Biobank traits, with increased prospects for biological interpretation (see Supplementary Table 13). We achieved this by using a multi-faceted functional enrichment model that includes coding, conserved, regulatory and LD-related annotations^8,12^.

Previous studies that assumed sparse genetic architectures achieved 3–5% increases in association power^17,20^. In detail, ref.^17^ reported a 5.0% increase in power (average *N*=57K for 18 traits) and ref.^20^ reported a 2.7% increase in power (*P* < 1 × 10^−8^; median *N_eff_* = 4/(1/*N_case_* + 1/*N_control_*) = 6K for 123 binary traits, median *N*=23K for 96 quantitative traits). (Ref.^20^ also reported a 13.7% increase in the number of ‘‘unsettled” associations (1 × 10^−10^ < P < 1 × 10^−8^), a metric that yields much larger increases.) In contrast, our polygenic approach achieved a 7% increase in association power (or 13% increase in power averaged across traits) in the interim UK Biobank analysis despite the larger sample size analyzed (average N=130K), which corresponds to smaller increases in power (see Figure 3). Ideally, we would have assessed those previous methods in the current study; however, we were unable to do so, either because no software implementation was available^20^, or because the available output (Bayes factors and posterior probabilities of association) was not directly comparable to the p-value thresholds used to assess significance in our study (and most GWAS studies)^17–19,21^. We instead elected to assess previous methods that could incorporate information from our polygenic functional enrichment model and produce p-value thresholds for hypothesis testing: Stratified FDR (S-FDR)^22^, Grouped Benjamini Hochberg (GBH)^23^, and Independent Hypothesis Weighting (IHW)^24^.

Stratifying SNPs based on predicted (tagged) variance was previously proposed by ref.^16^ (incorporating 10 functional annotations), which made a key contribution to the literature by highlighting the potential of this approach. The study demonstrated that this criteria improved replication rates, and also reported that it increased power when applying S-FDR^22^. However, S-FDR did not achieve proper null calibration in our simulations, even under random stratification, perhaps because S-FDR requires additional adjustments for the number of strata used ^29^. Furthermore, S-FDR, GBH, and IHW were all unable to correctly control false positives when LD-dependent stratification criteria (LDscore or BaseLD) were employed; as noted above, theoretical guarantees about false positives are unlikely to be valid when the stratification criteria incorporate the dependence structure between hypothesis tests^24^. Our approach bears some similarity to the multi-threshold association tests proposed by ref. ^14,15^, which use knowledge of the true effect size distribution to solve a convex optimization problem to determine appropriate thresholds. Given knowledge of the true effect size distribution, this approach is theoretically optimal^13,14^; however, this information is rarely available in practice and must be fixed a priori or approximated from the data^13–15^. Finally, although we employ a fundamentally different weighting strategy, our method draws on insights from ref. ^13^, which established the theoretical basis for data-driven p-value weighting.

We conclude with several limitations of our work. First, previous studies have demonstrated that complex traits often exhibit cell-type specific functional enrichments^4–11,17,32,33^, which we did not incorporate in this study. Incorporating cell-type-specific functional enrichments may further increase power, although care will be required to avoid overfitting since identifying critical cell types requires extensive model selection. Second, our modeling of MAF-dependent architectures is limited; while our baselineLD functional model includes MAF-bin annotations for common SNPs (MAF > 5%), it does not model MAF-dependent architectures for rare and low-frequency variants. A possible future direction would be to incorporate MAF-dependent annotations, e.g., via the widely used *α* model^34–36^. Third, we anticipate that GWAS will grow larger and more powerful in the years ahead, but the relative improvement of our method decreases as a function of absolute power. However, we anticipate that our method will continue to produce large relative improvements for disease phenotypes (as in Table 1), for which the ongoing challenge of recruiting disease cases will continue to limit effective sample size. Fourth, our UK Biobank replication of novel loci from the interim UK Biobank release could in principle be inflated by relatedness within the UK Biobank; however, our non-UK Biobank replication produced a concordant replication slope, suggesting that this effect is limited. Fifth, we evaluated our method only using European-ancestry samples. Although our previous work has provided evidence that functional enrichment is consistent across populations^10,37^, generalizing our results to non-European samples is currently an open question, as it is unclear whether functional enrichments inferred in large European samples should be incorporated. Despite these limitations, we anticipate that FINDOR will be a valuable and practical tool for leveraging polygenic functional enrichment to improve GWAS power.

## URLs

Open-source FINDOR software will be made publicly available prior to publication at https://github.com/gkichaev.

LDscore regression software: https://github.com/bulik/ldsc

LDscores for baselineLD model: https://data.broadinstitute.org/alkesgroup/LDSCORE/

UK Biobank Resource: http://www.ukbiobank.ac.uk/

BOLT-LMM v2.3 software http://data.broadinstitute.org/alkesgroup/BOLT-LMM/

BOLT-LMM association statistics (459K) http://data.broadinstitute.org/alkesgroup/UKBB/

## Acknowledgements

We are grateful to Yakir Reshef, Farhad Hormorzdiari, and Ruth Johnson for helpful discussions. This research was funded by NIH grants U01 HG009379, R01 MH101244, R01 MH107649 and R01 HG009120. This research was conducted using the UK Biobank resource under Application 16549.

## Online Methods

### FINDOR method

The aim of our method is to re-weight SNPs according to how well they tag heritability enriched categories. This is accomplished in two steps. First, we estimate a function that predicts the χ^2^ statistic (i.e. tagged variance) at each SNP using a comprehensive assortment of functional annotations which include coding, conserved and regulatory annoations^8^, as well as LD-dependent annotations^12^. The stratified LD score regression^8,12^ framework is a natural choice for this task. In stratified LD score regression, the association statistic at SNP j measured (or imputed) in *N_j_* individuals is expressed in terms of its tagging of studied annotations. Specifically,

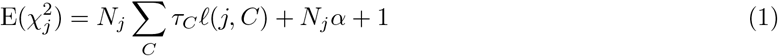

where *α* represents confounding biases^38^, *τ_C_* is the effect size on per-SNP heritability of annotation *C*, and *ℓ*(*j, C*) is the LD score which indicates the degree to which SNP *j* tags annotation *C*:

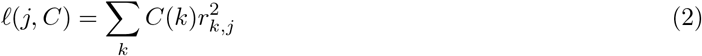

Here, *C*(*k*) is the value of annotation *C* at SNP *k* and 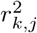 signifies the squared Pearson correlation coefficient between SNPs *k* and *j*^8,12^ (computed from 503 European individuals of the 1000 Genomes (V3) reference panel^39^). In a typical analysis, the quantity of interest is an estimate of 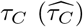 which can be interpreted as the strength of enrichment (or depletion) of heritability within annotation *C*. These values are obtained through a multivariate (weighted) regression of the observed χ^2^ statistics at HapMap3 SNPs against the corresponding values of *ℓ*(*j,C*). In this work, we use 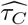 to predict the expected χ^2^ statistic at all GWAS SNPs. For a given SNP *j*, we have:

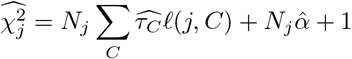

The 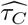 parameters can either be global estimates that are learned from the entire GWAS data set (restricted to HapMap3 SNPs), or chromosome-specific estimates that are learned from the remaining off-chromsome data. Empirically, we find that using the entire genome does not introduce false positives (see Figure 1).

Next, we stratify SNPs based on their expected χ^2^ into *B* distinct, evenly-sized bins. In practice, to ensure a sufficiently coarse partitioning of the data we set *B* = 100. For densely imputed data such as the UK Biobank this results in each bin *b* containing ≈ 100K SNPs. We then estimate the proportion of null 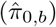 and alternate SNPs 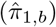 by fitting a cubic spline to the histogram of p-values as proposed by ref.^28^ and implemented in the q-value package^40^. Following ref.^23^ we weight each p-value by dividing the nominal p-value by the ratio of 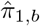 to 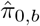. Intuitively, bins with higher proportion of true alternates will have their p-value weighted downward (i.e. made more significant). However, unlike ref.^23^, we normalize these weights to have mean one:

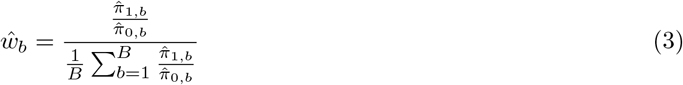

Theory developed in ref.^13^ suggests that despite the fact that 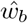 is learned in a data-dependent manner, a weighting scheme with this property preserves control of type I error since the number of weights we learn (i.e. 100) is significantly less than number of hypothesis test we perform.

### S-FDR, GBH and IHW methods

We adapted three previously proposed methodologies that leverage prior information to serve as comparators to our approach: Stratified False Discovery Rate (S-FDR)^22^, Grouped Benjamani Hochberg (GBH)^23^, and Independent Hypothesis Weighting (IHW)^24^. Because these are FDR-controlling procedures, we calibrate the expected level of FDR control required to match the more traditional criteria for genome-wide significance (*p* ≤ 5 × 10^−8^). We refer to this level of genome-wide FDR control as *q_GW_*, which we estimate as the maximum q-value^28^ amongst SNPs with p-values ≤ 5 × 10^−8^. We implemented S-FDR by binning SNPs according to various criteria used in this study. We then computed q-values for each bin and rejected all SNP within the bin whose q-value was less than *q_GW_*. This stratified FDR strategy is similar to Schork et al.^16^. GBH and IHW were implemented in the IHW (v1.1.3) and IHWpaper (v1.0.2) packages^24^ which we ran using the default setting and specified the level of FDR control to be *q_GW_*. GBH takes as input group labels which were identical to the groupings used with FINDOR and S-FDR, while IHW handled raw measurements of the auxiliary information (e.g., each SNP had its own unique value of predicted tagged variance under BaseLD).

### Functional Annotations

We employed the 75 functional annotations of the baselineLD model, which were previously demonstrated to be enriched for heritability across a wide variety of complex traits^12^ (see Supplementary Table 1). For clarity, we provide a brief description of the model’s contents below. This model is an extension of the 53 annotation baseline model developed in ref. ^8^. Briefly, the initial baseline model consisted of 24 main annotations to which 500bp flanking windows were added to create secondary annotations. These include histone modifications H3K4me1, H3K4me3, H3K4ac, H3K9ac, and H3K27ac that span multiple cell types; genic elements describing coding, 3’ UTR, 5’ UTR, promoter, and intronic regions; combined chromeHMM and Segway segmentations (7 states); Digital genomic footprint and transcription factor binding sites; DNase Hypersensitivity I sites; Super enhancers and FANTOM5 enhancers; and sites conserved across mammals (see ref.^8^ and references therein). The baseline model was augmented in ref.^12^ by adding four more binary annotations based on super-enhancers and typical enhancers, as well as two conserved annotations based on GERP++ scores. The baselineLD model was then created by adding ten common MAF bin annotations and six LD-related annotations (predicted allele age, LLD-AFR, recombination rate, nucleotide diversity, background selection statistic, and CpG content).

### Simulations

Simulations were based on real imputed genotypes of British ancestry individuals from the UK Biobank interim release (*N* =113K). We removed poorly imputed SNPs whose INFO score was less than 0.6, filtered out rare variants whose minor allele count was less than five in European individuals of the 1000 genomes, and additionally excluded the MHC region on chromosome six. This resulted in 9.6M SNPs for analysis. We randomly subsampled *N* individuals from this data set (in our main simulations, *N* =100K) and simulated continuous phenotypes under a polygenic model with normally distributed causal effect sizes and a specified number of causal variants. Genotypes were standardized so that each causal variant explained an equal proportion of the phenotypic variance. To induce functional enrichment, we altered the prior probability that a SNP was selected to be causal, setting this to be proportional to Var(*β_j_*) = Σ_*C*_ *C*(*j*)*τ_C_*. Empirically estimated enrichment parameters (*τ*’s) were obtained from a meta-analysis of the 31 traits reported in ref.^12^ (see Supplementary Table 1). This allowed our simulations to more closely reflect the complex, multi-faceted genetic architectures observed in real data. We note that functional enrichment was estimated without knowledge of the true functional enrichment used to simulate phenotypes. To obtain the baseLD-LOCO criteria, we estimated chromosome specific *τ*′s using off-chromosome data. Finally, we used PLINK v1.941 to compute association statistics for each SNP. The primary metric of interest in both real and simulated data was the number of independent GWAS hits (at a level of *p* < 5 × 10^−8^) that the various methodologies identified. We conservatively define independent hits using PLINK’s clumping algorithm with 5MB window and an *r*^2^ threshold of 0.01. Reference LD for this procedure was based on the same 113K British ancestry individuals for both simulations and real data analysis. To avoid over-counting loci where allelic heterogeneity was likely present in real data, we collapsed independent signals that were within 100KB of one another into a single locus.

### UK Biobank data set

We used BOLT-LMM ^27,30^ to compute mixed model association statistics. A key advantage of this approach that it allowed us to retain related individuals in this dataset, thereby maximizing power and data usage^27^. We performed basic QC on each trait following standard GWAS practices (see ref.^27^ for details). For each phenotype, we generated three sets of summary statistics based on individuals of self-reported European ancestry. The first set of summary statistics consisted of 145K individuals from the interim UK Biobank release^25,30^. This served as our “discovery” dataset and had mean sample size of « 130K across 27 independent traits (see below). We then created two additional sets of summary statistics derived from the full UKBiobank release^26^. Our “replication” dataset consisted of 314K individuals in the final release that were not present in the interim release (mean sample size = 283K). This dataset was used to verify findings in the discovery sample. Our “full” dataset was the entire compendium of 459K individuals (average *N* = 416K). While we computed summary statistics at 20 million SNPs which passed filtering and QC thresholds (see ref. ^27^), to ensure compatibility with simulations, we ran association analyses restricting to the same set of well-imputed ≈ 9.6M biallelic SNPs which, upon intersection, resulted in 9.6M SNPs for the interim release and 8.9M in the full release.

To avoid over-representation of certain phenotypic classes in our real data analysis that may bias our results, we constructed a set of 27 (roughly) independent and heritable traits, only retaining traits that exhibited a phenotypic correlation *r*^2^ < 0.1. To ensure adequate power to estimate functional enrichment, we also required that the traits have a heritability Z-score that was greater six in the 145K dataset to be included in our analysis^8^. An overview of the phenotypes analyzed in this work can be found in Table 1.

### Independent Non-UK Biobank data

To confirm the robustness of our findings we sought to replicate them in non-UK Biobank GWAS. We were able to obtain publicly available GWAS summary statistics for nine GWAS traits that were part of the 27 trait analysis (see Supplementary Table 10). As SNP coverage was not uniform, we intersected the data sets and only examined significant findings that were present both GWAS. When per-SNP sample sizes were unavailable, we used the max *N* obtained from the corresponding publication (see Supplementary Table 10). External GWAS alleles were polarized to the UK Biobank and standardized effect sizes were compared 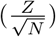.

### Replication Analysis

We carried out replication analysis in independent UK Biobank (27 traits; 3307 loci) and non-UK Biobank data (9 traits; 446 loci). To ensure compatibility across all traits and data sets, standardized effect sizes were computed by dividing Z-scores by the square root of the study sample size. To quantify replication, we computed the replication slope, defined as the slope resulting from a regression of the standardized effect sizes in the replication data versus the discovery data. We restricted our analysis to lead SNPs at independent, genome-wide significant loci in the discovery data that were also present in the replication data. We defined three class of loci: those that were genome-wide significant only using the Unweighted approach, only using FINDOR p-values, or using both methods. Because re-weighting could result in different lead SNPs at the same locus, we designated a locus as genome-wide significant using both methods if the lead SNP discovered by unweighted p-values had an *r*^2^ > 0.01 with the lead SNP discovered by FINDOR.

